# Activity-based anorexia enhances glutamatergic synaptic transmission and neuronal excitability within the nucleus accumbens of female mice

**DOI:** 10.64898/2026.02.12.705583

**Authors:** Lydia G. Bailey, Connor W. Christensen, Samantha E. Weed, Mohammed Moinul Islam, Amit Thakar, Jaedyn B. Brown, Shane T. Hentges, Travis E. Brown

## Abstract

Anorexia nervosa is a severe psychiatric disorder characterized by persistent food restriction and often excessive physical activity, implicating dysfunction in neural circuits governing motivation, reward, and behavioral persistence. The nucleus accumbens (NAc) is a central component of these circuits, yet synaptic and cellular adaptations within this region during anorexia-like states remain poorly defined. Using the activity-based anorexia (ABA) paradigm in adult female mice, we examined glutamatergic signaling and intrinsic neuronal properties in the NAc shell. ABA exposure produced rapid weight loss, reduced food intake, and progressively increased running-wheel activity. Biochemical analyses of NAc shell tissue revealed elevated membrane-associated GluA2 AMPA receptor protein. Consistent with this finding, whole-cell patch-clamp recordings from medium spiny neurons showed increased amplitude of spontaneous excitatory postsynaptic currents. ABA also enhanced intrinsic neuronal excitability, reflected by greater firing in response to depolarizing current injections. Together, these convergent biochemical and electrophysiological results demonstrate that ABA induces coordinated postsynaptic strengthening and increased intrinsic excitability in NAc shell medium spiny neurons. These adaptations suggest a sustained increase in accumbal output that may bias motivational circuit function and contribute to excessive activity and suppressed feeding during anorexia-like conditions, paralleling glutamatergic plasticity observed in other compulsive disorders, including substance use disorder.

## Introduction

Anorexia nervosa (AN) is a devastating psychiatric disorder with a lifetime prevalence of approximately 1%, and nearly the highest mortality rate of any psychiatric disorder (Meczekalski et al., 2013). Despite its prevalence and severity, effective and enduring treatments remain elusive, largely because the neurobiological mechanisms underlying AN are still poorly understood. Beyond severe caloric restriction, nearly 80% of individuals with AN engage in excessive physical activity, a paradoxical behavioral pattern that persists despite extreme energy deficits (reviewed in Ioannidis et al., 2021). This hyperactivity in the face of caloric deficit, coupled with the frequent comorbidity with substance use disorders (SUDs), major depressive disorder, anxiety and obsessive-compulsive disorders (Jordan et al., 2008), suggest an underlying dysregulation of the brain’s reward system in AN. Supporting this idea, numerous studies have reported altered reward-related behaviors (Ehrlich et al., 2015; Jappe et al., 2011; Keating et al., 2012; Wagner et al., 2007), as well as changes in neurotransmitter levels, and synaptic function in AN (Gorrell et al., 2020; Kaye et al., 2009; Nunn et al., 2012; Tharwani et al., 2025). Yet, despite these observations, a unifying model that explains how reward system dysfunction may drive the core symptoms of AN has yet to be established.

The overlap between AN and SUDs extends beyond clinical comorbidity to shared patterns of maladaptive behavior and reward dysfunction, which can interfere with daily functioning (Barbarich-Marsteller et al., 2011; Godlewska et al., 2017; O’Hara et al., 2015; Quintero, 2013). Neuroimaging studies reveal convergent abnormalities in key reward-related brain regions, including the nucleus accumbens (NAc), a critical hub for integrating motivational and hedonic signals. Further, interventions that target this circuitry, such as deep brain stimulation of the NAc, have shown early promise for alleviating symptoms of both AN and SUD (Campos-Fajardo et al., 2025; Ge et al., 2025) Finally, complimentary evidence from preclinical animal models further implicates glutamate signaling within the NAc as a key modulator of both drug intake and the aberrant feeding behaviors observed in AN, suggesting a shared circuit-level mechanism driving compulsive motivational states across these disorders.

An emerging body of work in animal models suggests a causative role for altered glutamate signaling within the NAc driving anorexia-like behaviors. In the activity-based anorexia (ABA) paradigm, rodents given continuous access to a running wheel, but restricted access to food, develop a maladaptive increase in running despite severe caloric deficit and progressive weight loss. However, when glutamatergic inputs from the prefrontal cortex (PFC) to the NAc shell were chemogenetically inhibited during the ABA paradigm, the phenotype was absent (Milton et al., 2021), showing that cortico-accumbens glutamate transmission is a key driver of running and weight loss in ABA. Consistent with this, exposure to ABA alters glutamatergic receptor expression within the NAc of rats exposed to ABA (Mottarlini et al., 2020). These preclinical findings align with clinical indications that interfering with glutamatergic signaling broadly has therapeutic promise in patients with AN. For example, pharmacological interventions that restore glutamatergic transmission, such as ketamine and similar compounds, and neuromodulatory approaches (Hermens et al., 2020; Ragnhildstveit et al., 2022) including deep brain stimulation of the NAc alleviate symptoms of AN (Campos-Fajardo et al., 2025), highlighting a central role for the NAc in mediating some of the symptoms of AN. Together, these lines of evidence suggest that dissecting how accumbal glutamate signaling is altered in ABA may both clarify the circuit mechanisms underlying AN-like behaviors and reveal new targets for intervention in patients.

In the current study, we used the ABA paradigm in adult female mice to identify how energy deficit and hyperactivity reshape glutamatergic signaling in the NAc shell. We found a selective enrichment of GluA2 protein surface expression, accompanied by strengthened AMPAR-mediated synaptic currents within the NAc shell. Additionally, we saw heightened intrinsic excitability of medium spiny neurons. Together, these convergent adaptations point to a NAc that is unusually responsive, reminiscent of the findings seen in models of SUD (D’Souza, 2015). By tracing these parallels, the present study frames anorexia-related behaviors not as anomalies of appetite, but as manifestations of enduring plasticity within reward circuitry, plasticity that persists, adapts, and ultimately compels. Hence, this work opens new avenues for probing the enduring neurobiological changes in reward signaling that may contribute to the manifestation and enduring nature of anorexia-like behaviors.

## Methods

### Ethical Approval

All experiments were approved by the Washington State University Animal Care and Use Committee (ASAF 7099) and were performed in accordance with the National Institutes of Health *Guide for the Care and Use of Laboratory Animals*, with efforts made to minimize the number of animals used and to minimize pain and suffering.

### Animals

Female C57BL/6J mice were obtained from the Jackson Laboratory (Bar Harbor, ME) and were housed on a phase-shifted (lights off at 10:00) 12-h light/dark cycle with temperature (20-22°C) and humidity held constant. Water was provided *ad libitum* throughout the study. Standard rodent chow (Product #5001; Animal Specialties, Quakertown, PA) was available *ad libitum* until initiation of the ABA paradigm. Mice were randomly assigned to experimental groups. At the start of the paradigm (ABA Day 0), mice were approximately 8 weeks old.

### Activity-based Anorexia Paradigm

Following arrival, mice were group-housed for 10 days to habituate to the facility and the phase-shifted light/dark cycle. Mice were subsequently single housed in experimental cages equipped with running wheels (Catalog #0297; Columbus Instruments, Columbus, OH). Animals were allowed to acclimate to the cages for 3 days prior to the start of a 5-day baseline period.

Mice were randomly assigned to one of four conditions: Activity-Based Anorexia (ABA), Food Restricted (FR), Exercise (EXE), or Sedentary Control (SED). Running wheels for SED and FR animals remained locked throughout the experiment. For ABA and EXE animals, running-wheel activity was recorded continuously in 15-min bins using Multi-Device Interface Software (Columbus instruments). Initial activity data during acclimation was analyzed to confirm proper entrainment to the light/dark cycle.

During the 5-day baseline, all animals had *ad libitum* access to food and water. Body weight and food intake were recorded daily 1 h prior to lights-out. Following baseline, food was removed from ABA and FR animals 2 h into the dark cycle (ABA day 0). On subsequent days (ABA day 1-6), ABA and FR animals received standard chow for a 2 h window beginning at the onset of the dark cycle. Food-anticipatory activity (FAA) was defined as running-wheel activity in the 4 h preceding food presentation. Body weight and food intake continued to be recorded for all animals 1 h prior to lights-out. Animals were removed from the study when they reached 80% of their baseline body weight or upon completion of day 6.

### Western Blots

On the day of removal from the paradigm, mice were deeply anesthetized with isoflurane, and the brains were immediately collected into ice-cold artificial cerebral spinal fluid (aCSF) containing (in mM) 124 NaCl, 2.5 KCl, 1.2 NaH_2_PO_4_, 24 NaHCO_3,_ 5 HEPES, 12.5 glucose, 2 MgCl_2_.7H2O, and 2 CaCl_2_.2H2O). 1 mm-thick sections were prepared on a Leica VT1000S vibratome in ice-cold aCSF and punches containing the nucleus accumbens shell were taken using a 1.25 mm diameter punch (Stoelting # 57401). The punches were then homogenized using a Dounce homogenizer in cold buffer (0.32 M sucrose containing 1 mM HEPES, 0.1 mM EGTA, 0.1 mM PMSF, pH = 7.4) containing protease and phosphatase inhibitors cocktail (ThermoScientific #78442).

Membrane fractions were prepared as previously described (Fumagalli et al., 2009) and were resuspended in buffer (20 mM HEPES, 0.1 mM DTT, 0.1 mM EGTA) with protease and phosphatase inhibitors (ThermoScientific #78442). Protein samples (10 µg) were mixed with Laemmli’s buffer containing β-mercaptoethanol and incubated at 99°C for 10 min and were separated via Polyacrylamide Gel Electrophoresis (PAGE) using 4–20 gradient gels (Bio-Rad # 4561095). After the separation, protein samples were transferred to PVDF membrane and blocked with 5% nonfat dry milk in Tris Buffered Saline with 0.1% Tween-20 and then incubated with primary antibody. GluA1 and GluA2 were probed separately on different membranes using rabbit anti-GluA1 (Invitrogen # PA5-32425, 1:500) or rabbit anti-GluA2 (Sigma Millipore # ZRB1008, 1:1000) primary antibodies and incubated overnight at 4°C. Membranes were then washed and incubated for 1 h at room temperature with horseradish peroxidase conjugated secondary antibody (goat anti-rabbit, Invitrogen # 656120, 1:10,000). The protein bands were visualized using freshly prepared Super-Signal West Femto Kit (Pierce) following the manufacturer’s instructions. A Gel Doc XRS digital imaging system (Bio-Rad) was used to capture images. Once imaged, the membranes were cut at the 75 kDa protein marker and the lower portion of the membranes were further probed for β-Actin. Membranes were incubated with HRP-conjugated mouse anti β-Actin (Invitrogen # MA5-15739-HRP) for 1 h at room temperature. Densitometry analysis was carried out using Image Lab software (Biorad). Protein levels of GluA1 and GluA2 were normalized to β-actin before calculating the GluA1/GluA2 ratio. Blots were compared by internally normalizing the blot to the corresponding SED control group.

### Whole-Cell Recordings

Once animals were removed from the behavioral paradigm, they were anesthetized using isoflurane and brains were extracted. Tissue was kept in a recovery solution containing (in mM): 93 NMDG, 2.5 KCl, 1.2 NaH2PO4, 30 NaHCO3, 20 HEPES, 25 Glucose, 5 sodium ascorbate, 2 thiourea, 3 sodium pyruvate, 10 MgSO4.7H20, and 0.5 CaCl2.2H20, and sliced on a Leica Microsystems vibratome at 300µm thickness. Slices were then incubated for 10 minutes in the same solution in a hot water bath, kept at ∼37°C. They were then transferred to a room temperature chamber for at least one hour in a solution containing (in mM): 92 NaCl, 2.5 KCl, 1.2 NaH2PO4, 30 NaHCO3, 20 HEPES, 25 glucose, 5 sodium ascorbate, 2 thiourea, 3 sodium pyruvate, 2 MgSO4.7H2O, and 2 CaCl2.2H2O.

Slices containing the NAc were placed in a recording chamber and continuously perfused with oxygenated aCSF at 30-33°C containing (in mM): 124 NaCl, 2.5 KCl, 1.2 NaH2PO4, 24 NaHCO3, 5 HEPES, 12.5 glucose, 2 MgSO4.7H2O, and 2 CaCl2.2H2O.

For spontaneous excitatory postsynaptic potential recordings, aCSF also contained 10µM bicuculline and 100 µM D-APV to block GABA and NMDA receptors, respectively. Patch pipettes (4-7 MΩ) were filled with (in mM): 130 KGluconate, 10 KCl, 1 EGTA, 10 HEPES, 0.6 NaGTP and 2 mgATP.

For excitability analysis, cells underwent current injection steps from -200 to 275 pA at 25 pA increments. Cells were discarded if they did not sustain firing throughout the protocol. For spontaneous EPSC recordings, cells were held at -70mV. Series access was assessed every minute, and cells were discarded if the series access changed more than 20% throughout the recording.

### Data analysis

Sample sizes were determined from our previous experiments and from related literature for electrophysiology experiments and behavior. For each electrophysiological measure, a maximum of three cells were recorded per animal to minimize oversampling. Prism 10.6.0 (GraphPad Software) was used to analyze the data. All data are presented as mean ± standard error of the mean. Differences were deemed significant with p<0.05.

Food anticipatory activity (FAA) was calculated using a paired t-test. Western blot data was calculated as the ratio of glutamate subunit to actin expression, then analyzed using a one-way ANOVA. We chose to analyze this data using a one-way ANOVA as opposed to a two-way in order to compare all groups to the control group to determine if any of the groups showed significant changes in protein expression. sEPSC event frequency and amplitude were calculated for all events, then 100 random events were selected for use in Kolmogorov-Smirnov test to ensure each cell was equally represented. Current-voltage relationship was analyzed using a mixed-effects analysis and Sidak’s multiple comparison test for post-hoc analysis. Intrinsic excitability was analyzed using a 2-way repeated measures ANOVA and Sidak’s multiple comparison test for post-hoc analysis. All remaining passive cell properties and action potential properties were analyzed using Welch’s t-tests.

## Results

Adult female mice exposed to the ABA paradigm (Figure 1A) showed rapid weight loss compared to age- and sex-matched sedentary control mice that did not undergo food restriction or have access to a running-wheel (SED, Figure 1B). No animals exposed to the ABA paradigm maintained more than 80% baseline bodyweight by day 6 of ABA exposure (Figure 1B, C). ABA animals exhibited increased running-wheel activity as the paradigm progressed (Figure 1D), which was observed to mainly be during the lights-on phase as food-anticipatory activity (FAA) which increased throughout the ABA paradigm (Figures 1E, F; t_22_ = 12.34; p<0.0001). In the few cases where there was a precipitous drop in running-wheel activity (Figure 1D), the animal reached the removal threshold (20% bodyweight loss) the next day, suggesting that physical condition, rather than reduced motivation, was likely responsible for the decline in running. While food intake remained steady across the days of the experiment for control animals, it decreased drastically and remained very low for animals in the ABA condition (Figure 1G). Altogether, these data confirm that the ABA model functioned as expected, with young adult female mice showing rapid weight loss and elevated running (Schalla & Stengel, 2019) that led all mice to reach threshold for removal within a few days.

**Figure 1.**
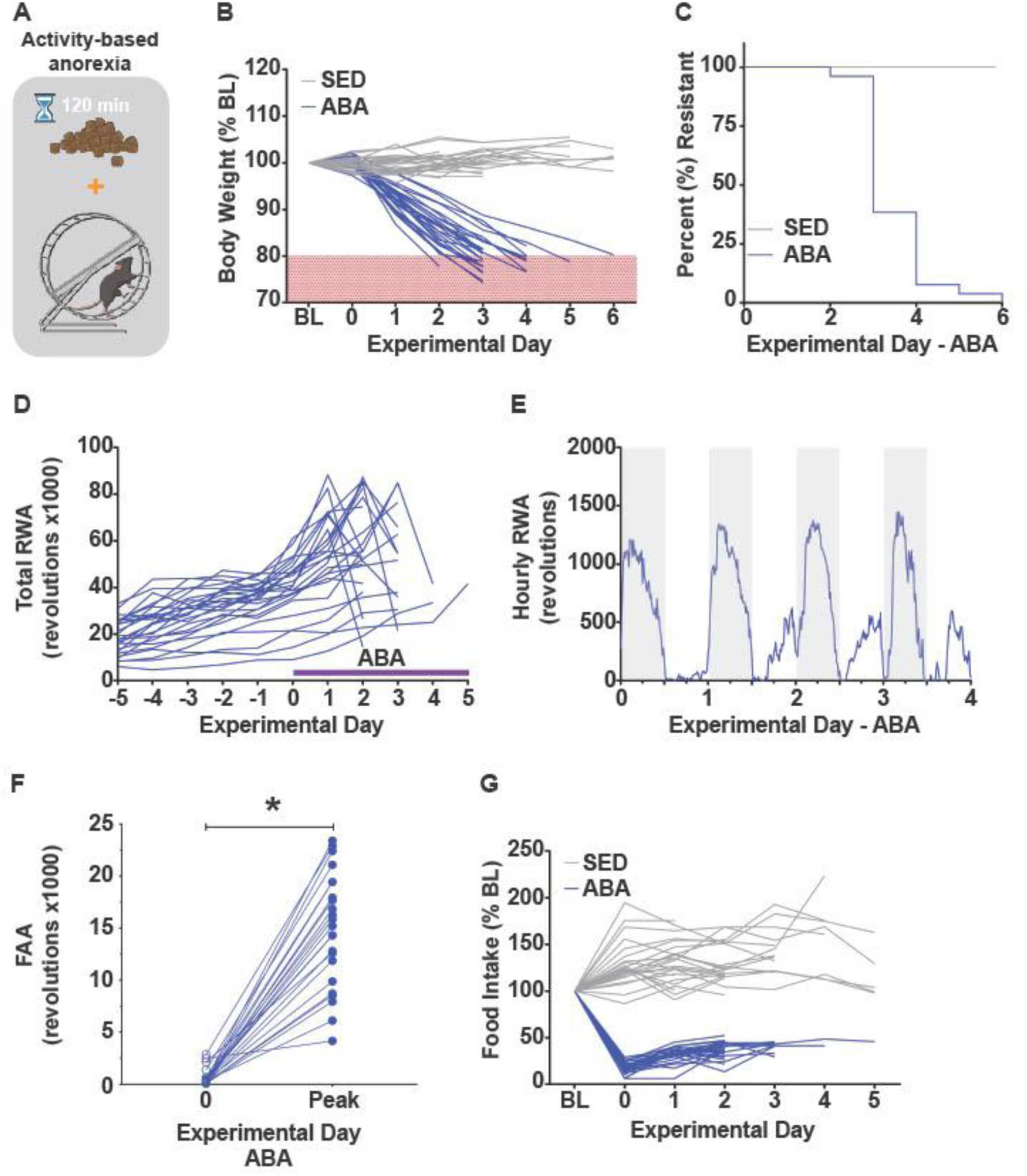
The ABA paradigm induces weight loss via decreased food consumption and increased running wheel activity. (A) A schematic illustrating that ABA animals were allowed 2 hours of food access and constant access to a running wheel. (B) Percent body weight of ABA (blue) and sedentary controls (grey) throughout ABA exposure, compared to their own average baseline (BL) body weight. (C) Survival curve for ABA and sedentary controls. Animals were removed from the study when daily weight measurement indicated the animal was at or below 80% of their baseline weight. (D) Total daily running-wheel activity for all ABA subjects. (E) Average running wheel activity by hour throughout the ABA paradigm. Shaded bars represent the animal’s dark phase, white bars represent the animal’s light phase. (F) Food anticipatory activity (FAA) on day 0 of paradigm versus the day of peak FAA for that animal. (G) Food intake for ABA and sedentary animals throughout the ABA paradigm normalized to their own average baseline food intake. N=25 mice SED, 26 mice ABA. Behavioral data here comes from all mice used throughout the studies such that a subset of the data here are also presented in figure 2. *Represents p<0.05.

A previous study revealed that there was altered expression of glutamate AMPA receptor subunits in the NAc of rats undergoing ABA (Mottarlini et al., 2020). To determine whether mice show a similar increase in calcium-permeable, GluA2 subunit-lacking AMPA receptors in the NAc as found in the rat, we analyzed GluA1 and GluA2 subunit expression in crude membrane fractions using Western blots. NAc membrane fractions prepared on the day that mice reached the removal threshold showed an increased GluA2 (one-way ANOVA, F =3.336, p=0.029; Dunnett’s control vs. ABA q_38_=2.685, p= 0.029) but not GluA1 (one-way ANOVA, p>0.5) expression compared to sedentary control mice (Figure 2A, B). Notably, no statistically significant changes in GluA1 or GluA2 expression were seen in animals exposed to food restriction or wheel running alone. Weight loss (Figure 2C), food intake (Figure 2D), and activity (Figures 2E, F) were as expected for the conditions of each group. This finding highlights the specificity of this neurological response to the combined ABA condition, as it was not induced by individual components of the model alone. The selective increase in GluA2 suggests that while rats and mice may both demonstrate increased glutamatergic signaling in the NAc during ABA, they do so via distinct mechanisms.

**Figure 2.**
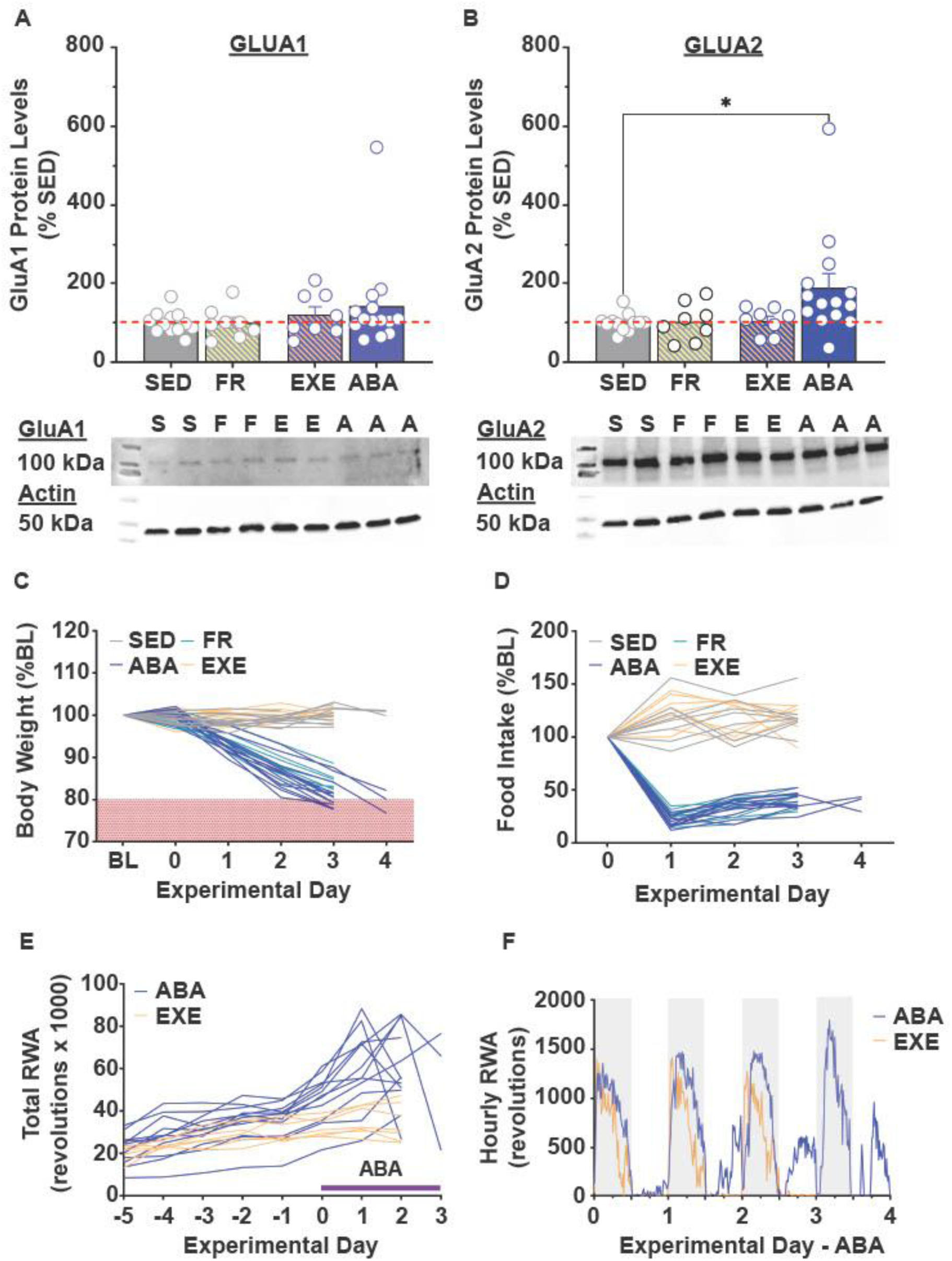
ABA increases GluA2 subunit expression. (A-B) GluA1 (A) and GluA2 (B) membrane protein levels in sedentary (SED; grey), food restricted (FR; yellow), exercise (EXE; pink), and ABA (blue) groups represented as percent change from sedentary (above) and representative western blots (below) of respective glutamate subunit (100kDa) and actin (47kDa) which was used for normalization of protein loading. Data points are calculated by the ratio of glutamate subunit to actin expression. (C-F) Body weight (C), food intake (D), total running activity (E) and hourly running activity (F) for all groups. *Represents p<0.05. C-F show behavioral data for the animals from which tissue was collected for the Western blots. N=12 SED, 8 FR, 8 EXE, 14 ABA.

We next sought to determine whether the observed change in protein expression correlated with detectable alterations in excitatory signaling in the NAc. To test this, we used whole-cell patch clamp electrophysiology and compared tissue from sedentary animals with tissue from ABA mice on the same day of removal from the paradigm. We focused on these two groups based on the protein data indicating that concomitant wheel-running and food restriction is necessary for a notable change in AMPAR expression. Overall changes in excitatory transmission within the NAc shell was assessed by examining spontaneous excitatory postsynaptic currents (sEPSC) onto medium spiny neurons (MSNs). In ABA animals sEPSCs exhibited greater amplitudes (Figure 3A; KS test, D=0.1848, p<0.0001) but no statistically significant change in event frequency (Figure 3B; KS test p>0.5). These data suggest an increase in postsynaptic glutamatergic signaling without an increase in the number of presynaptic release events.

**Figure 3.**
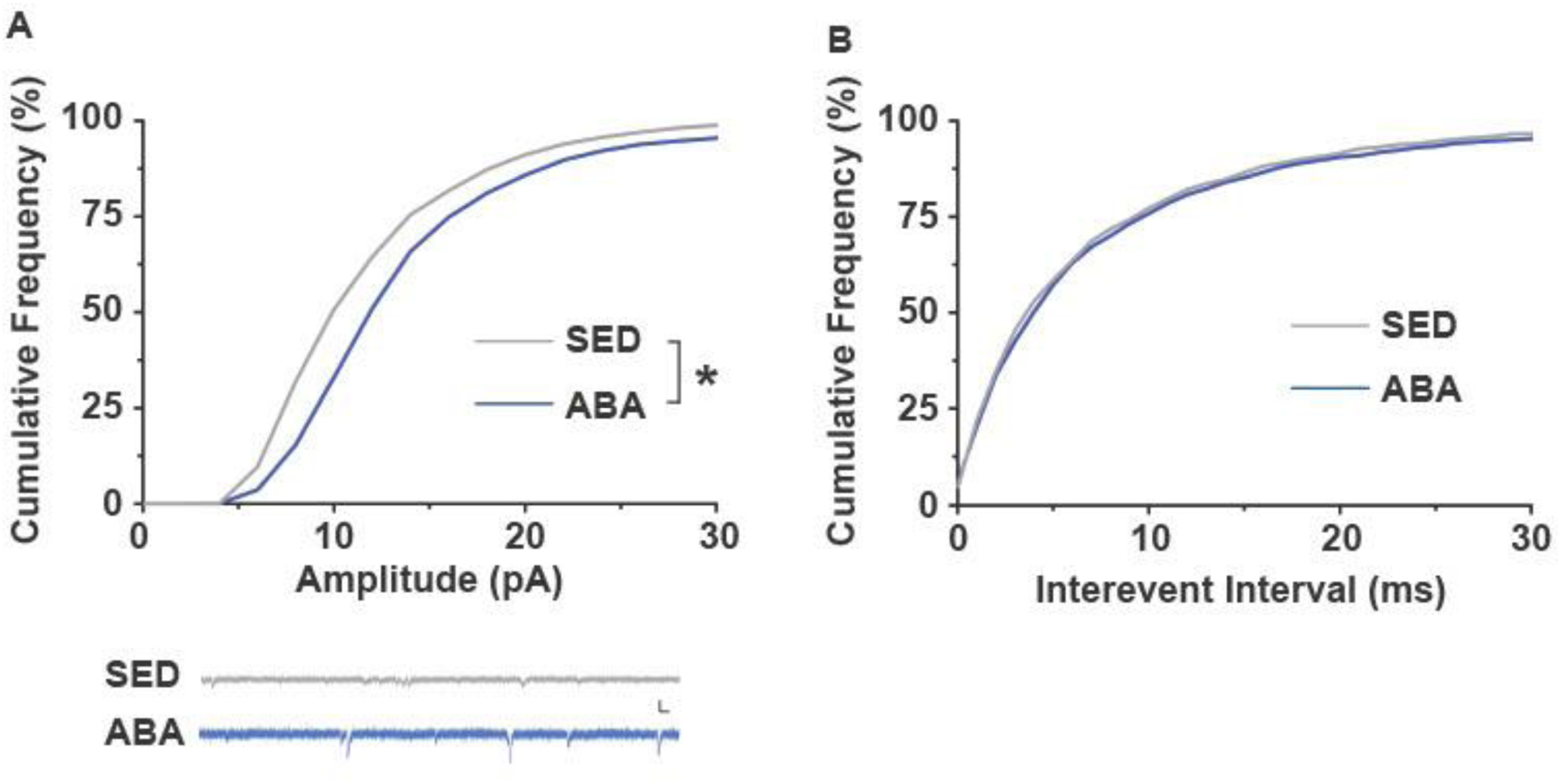
ABA increases sEPSC amplitude but not frequency. (A) Cumulative distribution of ABA (blue) and sedentary (grey) spontaneous excitatory postsynaptic current (sEPSC) amplitudes. (B) Cumulative distribution of ABA and sedentary sEPSC interevent intervals. Data for each curve represents the average of 100 random events from each cell. N=13-14 cells from 8-10 animals. *Indicates p<0.05

Reasoning that undergoing the ABA paradigm might change cellular properties of NAc shell MSNs, we examined passive membrane properties. ABA animals showed significantly increased conductance at negative current injection steps (Figure 4A; - 200pA to -75pA, all p<0.05), suggesting increased excitability. While there was no difference in resting membrane potential between control and ABA animals (Figure 4B), neurons from ABA animals exhibited increased membrane resistance (Figure 4C; Welch’s t_23.4_=4.509, p<0.001) and an increased time constant (Figure 4D; Welch’s t_18.2_ =2.404; p=0.027). However, the capacitance of cells from ABA animals compared to control were not statistically different (Figure 4E), indicating consistency of neuronal resting state between groups, but an increase in overall responsivity to excitatory currents.

**Figure 4.**
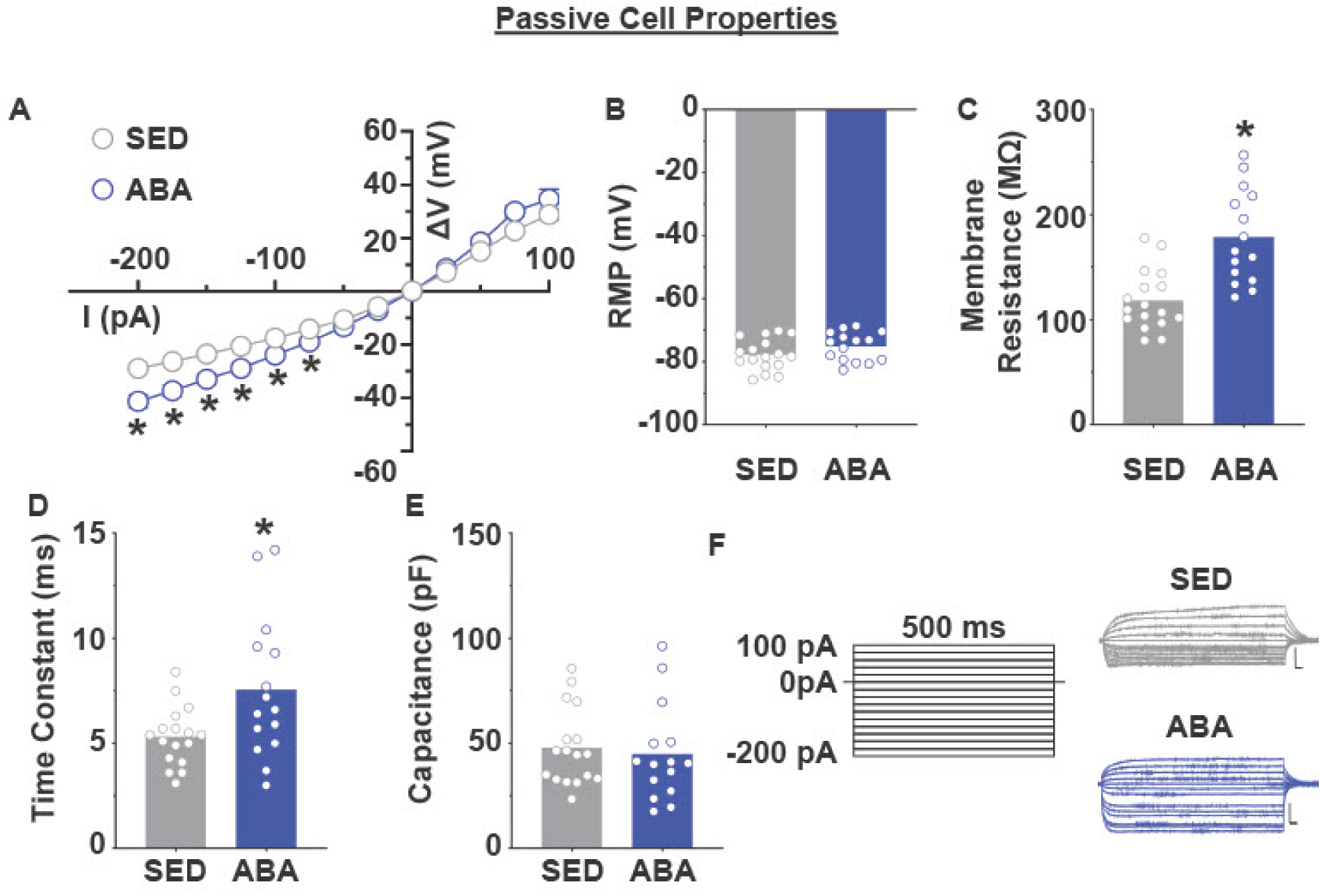
ABA alters intrinsic membrane properties of NAc shell neurons. (A) Current-voltage (I-V) relationship for neurons from sedentary (grey) and ABA (blue) animals. (B-E) Bar graphs representing resting membrane potential (B), membrane resistance (C), time constant (D), and capacitance (E) of sedentary controls and ABA treated animals. (F) Representative voltage responses to various current injection steps. N=16-19 cells from 7-8 animals. *Indicates p<0.05

Finally, we assessed intrinsic excitability and action potential properties of NAc shell MSNs. Cells from ABA animals demonstrated increased firing rate compared to sedentary controls (Figure 5A; 2-way RM ANOVA main effect of treatment, F_1, 33_ =8.15, p=0.007). While there was also a significant current x treatment interaction (F_1.521, 50.21_ = 5.076, p=0.016), post-hoc analyses did not reveal a statistically significant difference at any individual current step. In line with this, cells from ABA animals showed an associated decreased rheobase (Figure 5B; Welch’s t_27.53_ =2.444, p=0.021), suggesting that less current is required to elicit action potentials. Further, the after hyperpolarization (AHP) amplitude in the ABA group was significantly decreased compared to sedentary controls (Figure 5F; Welch’s t_21.61_ =2.293, p=0.031). Interestingly, the threshold, amplitude, and half-width showed no statistically significant changes from control (all p>0.2), demonstrating that the properties of the action potential itself remain largely unchanged.

**Figure 5.**
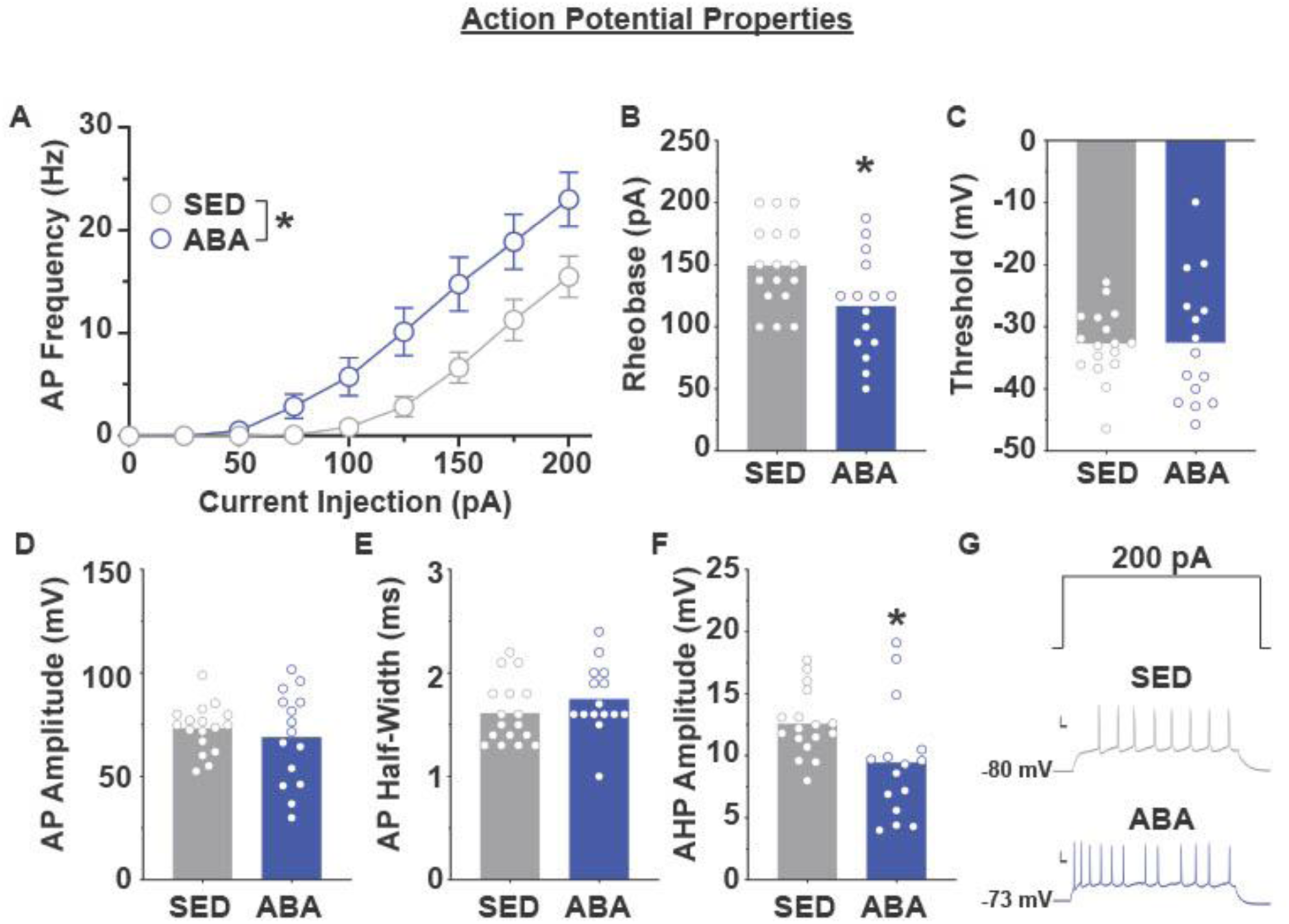
ABA increases neuronal excitability within the NAc shell. (A) Neuronal excitability at increasing current injections for cells from sedentary (grey) and ABA (blue) animals. (B-F) Action potential properties of neurons from sedentary and ABA groups showing the rheobase (B), action potential threshold (C), amplitude (D), half-width (E), and after hyperpolarization amplitude (F). (G) Representative traces showing neuronal excitability at 200 pA injected current. N=16-19 cells from 7-8 animals. *Indicates p<0.05.

## Discussion

In this study, we found that the activity-based anorexia (ABA) paradigm in mice induces synaptic plasticity within the NAc shell which leads to increased glutamatergic signaling. The observed changes are similar to those seen in models of substance use disorders, suggesting that enhanced excitatory signaling within the NAc shell may contribute to the disrupted reward circuitry and compulsive-like behavior observed in ABA. Further, these findings might provide some insight into why glutamatergic modulators (ketamine) and NAc-targeted neuromodulation (deep brain stimulation) show some promise in patients with AN, although this remains to be explored.

The initial observation of a statistically significant increase in GluA2, but not GluA1 protein levels in the membrane fraction of NAc shell neurons from ABA exposed mice compared to sedentary, food restricted, and exercise-only control mice was surprising. Based on prior work in rat, we expected a relative decrease in GluA2/increase in GluA1 (Mottarlini et al., 2020) but our results show a potential increase in calcium-impermeable AMPA receptors rather than enhanced calcium-permeable AMPARs. The enhancement of calcium-permeable GluA2-lacking AMPARs is generally considered a hallmark of initial synaptic plasticity (Man, 2011). In contrast, a lasting increase in GluA2-containing AMPARs as seen in the results here is more often associated with the maintenance of synapses following a period of plasticity (Man, 2011). This discrepancy between our mouse data and the published rat data could be explained by several factors. For example, it is possible that we missed an earlier, transient upregulation of GluA1 subunits, as we only assessed tissue at a single timepoint (threshold weight loss, ∼ day 3 of ABA). Additionally, there may be a genuine mechanistic difference between the two species, or slightly differing experimental protocols may induce distinct alterations. Finally, although changes in accumbens glutamatergic signaling are observed following both drug self-administration and palatable food exposure, data from Dingess et al. demonstrate that the patterns of AMPA/NMDA receptor–mediated plasticity elicited by high-fat food cues differ from those typically reported in drug models, highlighting that diet-induced glutamatergic adaptations do not precisely mirror the drug-evoked synaptic changes described in addiction studies (Dingess et al., 2017).

We next sought to determine whether these protein-level changes translated to functional alterations in glutamate signaling within the NAc shell MSNs. For these functional studies, we compared several cellular parameters from ABA animals (on their day of removal) with time-matched sedentary controls. We focused on these two groups because our protein data clearly showed that neither food restriction nor exercise alone was sufficient to alter GluA2 expression. This strongly suggests that it is the synergistic effect of food restriction and exercise that drives the observed changes in glutamate signaling in the NAc.

The electrophysiological results suggest that at least some of the increased GluA2 protein is in the form of functional tetrameric AMPARs within the membrane based on the functional increase in cell signaling and activity. The increase in spontaneous excitatory postsynaptic current (sEPSC) amplitude (Figure 3A) suggests there is a change on the postsynaptic membrane influencing receptor abundance or sensitivity, as is expected with an increase in AMPA receptor abundance throughout the region. The lack of significant changes in the frequency of sEPSCs (Figure 3B) suggest that the main influences seen on spontaneous excitatory events are not mediated by presynaptic influences on vesicular release, but increased responsivity of the postsynaptic cell. Further, ABA-induced calcium-impermeable AMPA receptor insertion into the membrane is supported by the passive membrane properties of cells recorded from the NAc shell. In addition to the AMPAR insertion mediated increase in glutamatergic activity, we found an increase in excitability of cells within the NAc shell. Cells from animals exposed to the ABA paradigm show increased membrane resistance and time constant (Figure 4), both markers of increased excitability of the cell. Further, increased neuronal excitability, decreased rheobase, and decreased after hyperpolarization (Figure 5) suggest overall increases in the excitability of cells within the NAc shell.

AMPA receptors within the NAc shell, while playing a role in reward-seeking behaviors, have also shown a specific role in mediating ingestive behavior due to the known projections stemming from this region (Maldonado-Irizarry et al., 1995). A coordinated reduction in excitatory projections to the NAc shell initiates food seeking (Reed et al., 2018), suggesting that overall changes in activity level of this region may contribute to disruption of natural food seeking behaviors. Previously, NAc shell D1 MSNs were shown to exert causal control over lateral hypothalamic inhibition, with activation of these projections resulting in cessation of food consumption regardless of hunger state (O’Connor et al., 2015). Further, connections from the NAc shell to the ventral tegmental area (VTA) have also been implicated in mediating food-seeking (Krause et al., 2010). Future studies are required to investigate the upstream and downstream regions that are involved in the altered glutamatergic circuit in ABA animals. However, we favor the hypothesis that PFC stemming projections converge on the NAc shell, altering overall motivational state within the reward circuitry pathway. Evidence for this hypothesis is provided by the findings of Milton et al. where they showed that inhibiting PFC→NAc glutamatergic inputs prevented dire weight loss in ABA in rats (Milton et al., 2021) which merges nicely with our observation of enhanced excitation in the NAc shell during ABA. The dysregulated glutamatergic signaling within the NAc may then increase excitatory output to target regions including the lateral hypothalamus. Additional projections may also be involved, explaining the divergent motivations for exercise compared to food. Regardless of the precise mechanism, the role of the NAc shell as a central hub for both homeostatic and hedonic reward seeking (Marinescu & Labouesse, 2024) suggests that altered signaling within this region may underlie some of the symptoms seen in AN patients.

In a paradoxical manner, reduced food intake in ABA and AN is accompanied by increased motivation for exercise (wheel-running activity). Inactivation of the NAc shell has been shown to reduce motivation to run (Basso & Morrell, 2015), suggesting that an overall increase in glutamatergic signaling within this region, as shown here, could reasonably result in increased motivation for running. Moreover, voluntary exercise has been shown to increase excitatory currents in D1, but not D2 expressing neurons, which was not present with forced exercise (Gan et al., 2023). This suggests that altered dopaminergic transmission may underlie some of the effects of the ABA paradigm and may create a feedback loop further enhancing motivation to run. The present studies did not target D1- or D2-expressing neurons, but the results are consistent with likely changes in excitability of D1-expressing cells, as co-activation of D1 and inactivation of D2 MSNs within the NAc shell have previously been shown to precipitate weight loss in anorexia models (Walle et al., 2024). However, future studies should investigate the roles of D1 and D2 expressing neurons individually to better understand their roles in AN.

Altogether, the present results suggest a modulatory role of glutamatergic signaling within the NAc shell for mediating the altered motivation observed when food-restriction and activity are paired, as occurs in the ABA model and in most cases of AN. The increase in calcium-impermeable AMPA receptors suggests a long-lasting, stable form of increased glutamatergic signaling throughout the NAc shell, ultimately contributing to altered motivation for various reward driven behaviors simultaneously.

The findings here may help explain why human patients with AN show benefits from manipulations to the glutamatergic system (Hermens et al., 2020) and the NAc (Campos-Fajardo et al., 2025). While the downstream targets of this altered glutamatergic signaling could include targets such as the lateral hypothalamus or the VTA, the exact projections involved remain to be determined. A limitation of the present study is that it did not include exercise-only or food-restricted-only control groups in the electrophysiology studies. The decision to focus on sedentary versus ABA animals was based on the protein results showing a change in glutamate receptor expression only when food restriction and running were paired. However, we cannot rule out the possibility that the electrophysiologic approach could be more sensitive and could detect a change in one or both of those groups that was not indicated by Western blot data. Future work should interrogate this possibility. Nonetheless, the NAc’s role as a central hub in maintaining homeostasis and influencing motivation in ABA and AN is becoming increasingly clear.

## Ethical use of animals

All experiments were approved by the Institutional Animal Care and Use Committees at Washington State University in accordance with the National Institute of Health’s Guide for the Care and Use of Laboratory Animals.

## Data Availability Statement

The datasets generated for this study are available upon request from the corresponding author.

## Acknowledgements

The authors gratefully acknowledge the staff of the Washington State University Animal Care Facility for their outstanding animal husbandry, veterinary support, and technical assistance.

## Author Contributions

All work was completed in the labs of T.E.B. and S.T.H. T.E.B. and S.T.H. designed experiments. L.G.B., C.W.C, S.E.W., M.M.I., and A.T. collected data. L.G.B., T.E.B., S.T.H., S.E.W., A.T., and J.B.B. analyzed data. L.G.B., S.T.H., T.E.B., A.T., and C.W.C. wrote the manuscript.

All authors have approved the final version of the manuscript and agree to be accountable for all aspects of the work.

## References

Barbarich-Marsteller, N. C., Foltin, R. W., & Walsh, B. T. (2011). Does Anorexia Nervosa Resemble an Addiction? Current Drug Abuse Reviews, 4(3), 197–200. 10.2174/1874473711104030197

Basso, J. C., & Morrell, J. I. (2015). The medial prefrontal cortex and nucleus accumbens mediate the motivation for voluntary wheel running in the rat. Behavioral Neuroscience, 129(4), 457–472. 10.1037/bne0000070

Campos-Fajardo, S., Lacouture-Silgado, I., Agudelo-Arrieta, M., & Zorro, Ó. F. (2025). Deep brain stimulation of the nucleus accumbens for the management of eating disorders: A scoping review. Clinical Neurology and Neurosurgery, 256, 109029. 10.1016/j.clineuro.2025.109029

Dingess, P. M., Darling, R. A., Derman, R. C., Wulff, S. S., Hunter, M. L., Ferrario, C. R., & Brown, T. E. (2017). Structural and Functional Plasticity within the Nucleus Accumbens and Prefrontal Cortex Associated with Time-Dependent Increases in Food Cue-Seeking Behavior. Neuropsychopharmacology, 42(12), 2354–2364. 10.1038/npp.2017.57

D’Souza, M. S. (2015). Glutamatergic transmission in drug reward: Implications for drug addiction. Frontiers in Neuroscience, 9. 10.3389/fnins.2015.00404

Ehrlich, S., Geisler, D., Ritschel, F., King, J. A., Seidel, M., Boehm, I., Breier, M., Clas, S., Weiss, J., Marxen, M., Smolka, M. N., Roessner, V., & Kroemer, N. B. (2015). Elevated cognitive control over reward processing in recovered female patients with anorexia nervosa. Journal of Psychiatry and Neuroscience, 40(5), 307–315. 10.1503/jpn.140249

Fumagalli, F., Pasini, M., Frasca, A., Drago, F., Racagni, G., & Riva, M. A. (2009). Prenatal stress alters glutamatergic system responsiveness in adult rat prefrontal cortex. Journal of Neurochemistry, 109(6), 1733–1744. 10.1111/j.1471-4159.2009.06088.x

Gan, Y., Dong, Y., Dai, S., Shi, H., Li, X., Wang, F., Fu, Y., & Dong, Y. (2023). The different cell-specific mechanisms of voluntary exercise and forced exercise in the nucleus accumbens. Neuropharmacology, 240, 109714. 10.1016/j.neuropharm.2023.109714

Ge, S., Wang, X., Chen, L., Li, N., Li, Y., Cai, Y., Wang, X., Li, W., Su, M., Zheng, Z., Li, J., Wang, X., Qiu, C., Wang, J., Liu, T., Qu, Y., & Gao, G. (2025). Effects of deep brain stimulation of the nucleus accumbens and anterior limb of the internal capsule on heroin addiction: Over five years of long-term follow-up in a prospective open-label pilot study. Translational Psychiatry, 15(1), 415. 10.1038/s41398-025-03635-6

Godlewska, B. R., Pike, A., Sharpley, A. L., Ayton, A., Park, R. J., Cowen, P. J., & Emir, U. E. (2017). Brain glutamate in anorexia nervosa: A magnetic resonance spectroscopy case control study at 7 Tesla. Psychopharmacology, 234(3), 421–426. 10.1007/s00213-016-4477-5

Gorrell, S., Collins, A. G. E., Le Grange, D., & Yang, T. T. (2020). Dopaminergic activity and exercise behavior in anorexia nervosa. OBM Neurobiology, 4(1), 10.21926/obm.neurobiol.2001053. 10.21926/obm.neurobiol.2001053

Hermens, D. F., Simcock, G., Dutton, M., Bouças, A. P., Can, A. T., Lilley, C., & Lagopoulos, J. (2020). Anorexia nervosa, zinc deficiency and the glutamate system: The ketamine option. Progress in Neuro-Psychopharmacology and Biological Psychiatry, 101, 109921. 10.1016/j.pnpbp.2020.109921

Ioannidis, K., Hook, R. W., Grant, J. E., Czabanowska, K., Roman-Urrestarazu, A., & Chamberlain, S. R. (2021). Eating disorders with over-exercise: A cross-sectional analysis of the mediational role of problematic usage of the internet in young people. Journal of Psychiatric Research, 132, 215–222. 10.1016/j.jpsychires.2020.11.004

Jappe, L. M., Frank, G. K. W., Shott, M. E., Rollin, M. D. H., Pryor, T., Hagman, J. O., Yang, T. T., & Davis, E. (2011). Heightened sensitivity to reward and punishment in anorexia nervosa. International Journal of Eating Disorders, 44(4), 317–324. 10.1002/eat.20815

Jordan, J., Joyce, P. R., Carter, F. A., Horn, J., McIntosh, V. V. W., Luty, S. E., McKenzie, J. M., Frampton, C. M. A., Mulder, R. T., & Bulik, C. M. (2008). Specific and nonspecific comorbidity in anorexia nervosa. International Journal of Eating Disorders, 41(1), 47–56. 10.1002/eat.20463

Kaye, W. H., Fudge, J. L., & Paulus, M. (2009). New insights into symptoms and neurocircuit function of anorexia nervosa. Nature Reviews Neuroscience, 10(8), 573–584. 10.1038/nrn2682

Keating, C., Tilbrook, A. J., Rossell, S. L., Enticott, P. G., & Fitzgerald, P. B. (2012). Reward processing in anorexia nervosa. Neuropsychologia, 50(5), 567–575. 10.1016/j.neuropsychologia.2012.01.036

Krause, M., German, P. W., Taha, S. A., & Fields, H. L. (2010). A Pause in Nucleus Accumbens Neuron Firing Is Required to Initiate and Maintain Feeding. Journal of Neuroscience, 30(13), 4746–4756. 10.1523/JNEUROSCI.0197-10.2010

Maldonado-Irizarry, C. S., Swanson, C. J., & Kelley, A. E. (1995). Glutamate receptors in the nucleus accumbens shell control feeding behavior via the lateral hypothalamus. Journal of Neuroscience, 15(10), 6779–6788. 10.1523/JNEUROSCI.15-10-06779.1995

Man, H.-Y. (2011). GluA2-lacking, calcium-permeable AMPA receptors—Inducers of plasticity? Current Opinion in Neurobiology, 21(2), 291–298. 10.1016/j.conb.2011.01.001

Marinescu, A.-M., & Labouesse, M. A. (2024). The nucleus accumbens shell: A neural hub at the interface of homeostatic and hedonic feeding. Frontiers in Neuroscience, 18. 10.3389/fnins.2024.1437210

Meczekalski, B., Podfigurna-Stopa, A., & Katulski, K. (2013). Long-term consequences of anorexia nervosa. Maturitas, 75(3), 215–220. 10.1016/j.maturitas.2013.04.014

Milton, L. K., Mirabella, P. N., Greaves, E., Spanswick, D. C., van den Buuse, M., Oldfield, B. J., & Foldi, C. J. (2021). Suppression of Corticostriatal Circuit Activity Improves Cognitive Flexibility and Prevents Body Weight Loss in Activity-Based Anorexia in Rats. Biological Psychiatry, 90(12), 819–828. 10.1016/j.biopsych.2020.06.022

Mottarlini, F., Bottan, G., Tarenzi, B., Colciago, A., Fumagalli, F., & Caffino, L. (2020). Activity-Based Anorexia Dynamically Dysregulates the Glutamatergic Synapse in the Nucleus Accumbens of Female Adolescent Rats. Nutrients, 12(12), 3661. 10.3390/nu12123661

Nunn, K., Frampton, I., & Lask, B. (2012). Anorexia nervosa – A noradrenergic dysregulation hypothesis. Medical Hypotheses, 78(5), 580–584. 10.1016/j.mehy.2012.01.033

O’Connor, E. C., Kremer, Y., Lefort, S., Harada, M., Pascoli, V., Rohner, C., & Lüscher, C. (2015). Accumbal D1R Neurons Projecting to Lateral Hypothalamus Authorize Feeding. Neuron, 88(3), 553–564. 10.1016/j.neuron.2015.09.038

O’Hara, C. B., Campbell, I. C., & Schmidt, U. (2015). A reward-centred model of anorexia nervosa: A focussed narrative review of the neurological and psychophysiological literature. Neuroscience & Biobehavioral Reviews, 52, 131–152. 10.1016/j.neubiorev.2015.02.012

Quintero, G. C. (2013). Role of nucleus accumbens glutamatergic plasticity in drug addiction. Neuropsychiatric Disease and Treatment, 9, 1499–1512. 10.2147/NDT.S45963

Ragnhildstveit, A., Slayton, M., Jackson, L. K., Brendle, M., Ahuja, S., Holle, W., Moore, C., Sollars, K., Seli, P., & Robison, R. (2022). Ketamine as a Novel Psychopharmacotherapy for Eating Disorders: Evidence and Future Directions. Brain Sciences, 12(3), 382. 10.3390/brainsci12030382

Reed, S. J., Lafferty, C. K., Mendoza, J. A., Yang, A. K., Davidson, T. J., Grosenick, L., Deisseroth, K., & Britt, J. P. (2018). Coordinated Reductions in Excitatory Input to the Nucleus Accumbens Underlie Food Consumption. Neuron, 99(6), 1260–1273.e4. 10.1016/j.neuron.2018.07.051

Schalla, M. A., & Stengel, A. (2019). Activity Based Anorexia as an Animal Model for Anorexia Nervosa–A Systematic Review. Frontiers in Nutrition, 6. 10.3389/fnut.2019.00069

Tharwani, Z. H., Shaeen, S. K., Zahid, K., Ahmed, S., Naqrashi, R., Murtaza, A., Mughal, S., Hasanain, M., Anjum, M. U., Eljack, M. M. F., & Zaidi, S. W. (2025). Exploring the link between dopamine dysregulation and eating disorders: A narrative review. *Journal of Neuroendocrinology*, *n/a*(n/a), e70070. 10.1111/jne.70070

Wagner, A., Aizenstein, H., Venkatraman, V. K., Fudge, J., May, J. C., Mazurkewicz, L., Frank, G. K., Bailer, U. F., Fischer, L., Nguyen, V., Carter, C., Putnam, K., & Kaye, W. H. (2007). Altered Reward Processing in Women Recovered From Anorexia Nervosa. American Journal of Psychiatry, 164(12), 1842–1849. 10.1176/appi.ajp.2007.07040575

Walle, R., Petitbon, A., Fois, G. R., Varin, C., Montalban, E., Hardt, L., Contini, A., Angelo, M. F., Potier, M., Ortole, R., Oummadi, A., De Smedt-Peyrusse, V., Adan, R. A., Giros, B., Chaouloff, F., Ferreira, G., de Kerchove d’Exaerde, A., Ducrocq, F., Georges, F., & Trifilieff, P. (2024). Nucleus accumbens D1- and D2-expressing neurons control the balance between feeding and activity-mediated energy expenditure. Nature Communications, 15(1), 2543. 10.1038/s41467-024-46874-9

